# RNG2 tethers the conoid to the apical polar ring in *Toxoplasma gondii*: a key mechanism in parasite motility and invasion

**DOI:** 10.1101/2024.08.29.610260

**Authors:** Romuald Haase, Bingjian Ren, Nicolas Dos Santos Pacheco, Rémy Visentin, Bohumil Maco, Ricardo Mondragon Flores, Oscar Vadas, Dominique Soldati-Favre

## Abstract

In *Toxoplasma gondii*, the conoid is a dynamic organelle composed of spiraling tubulin fibers that extrudes during egress, gliding motility, and invasion. This organelle traverses the apical polar ring (APR) in response to calcium waves and plays a critical role in controlling parasite motility. While the actomyosin-dependent extrusion of the conoid is beginning to be understood, the mechanism by which it is anchored apically to the APR remains unclear. RNG2, a protein localized at both the conoid and the APR, has emerged as a key candidate for this function. By combining iterative ultrastructure expansion microscopy and immunoelectron microscopy we discovered that RNG2 forms 22 tethers between the APR and the conoid. The unique biochemical properties of RNG2, including several proteolytic processing events and its ability to form concatenations, enable it to function as a dynamic bridge between these structures. Conditional depletion of RNG2 resulted in the conoid organelle detaching from the APR without compromising the integrity of its structure, thereby confirming RNG2 essential tethering role. Although microneme secretion remains normal, parasites lacking RNG2 were unable to move and impaired in rhoptry discharge, highlighting the conoid’s crucial role in parasite motility and invasion. RNG2 is a pivotal protein that ensures conoid functionality in *Coccidia*.

## Introduction

The phylum Apicomplexa comprises single-cell eukaryotic parasites of high medical relevance, most notably *Plasmodium* spp., *Cryptosporidium* spp., and *Toxoplasma gondii* [1]. Members of this phylum share an elaborate apical complex composed of specialized secretory organelles called rhoptries and micronemes, as well as tubulin-based cytoskeletal elements [2, 3]. The apical complex critically coordinates organelle secretion and activation of the actomyosin system to drive gliding motility, invasion, and egress from infected cells [4, 5].

The shape of *T. gondii* is determined by scaffolding components, including the inner membrane complex (IMC), the alveolin network and the subpellicular microtubules (SPMTs), which collectively provide rigidity to the parasite [6, 7] (Fig. 1). The 22 SPMTs extend approximately two-thirds of the parasite length and emerge from a protein-based structure called the apical polar ring (APR) [8, 9] (Fig. 1). The conoid consists of a cone of spiraling tubulin fibers [10], surmounted by two preconoidal rings (PCRs) serving as hubs for the assembly of the gliding machinery [11]. The inside of the conoid hosts two short intraconoidal microtubules (ICMTs) [10] with aligned microtubule-associated vesicles predicted to participate in rhoptry discharge [12–14] (Fig. 1). The conoid is a dynamic organelle that extrudes and retracts through the APR in response to changes in intracellular calcium levels [15, 16]. Conoid extrusion relies on F-actin produced by formin 1 (FRM1) positioned at the PCRs, and is powered by myosin H (MyoH), anchored to the cone [11]. The dynamics of the conoid serve as a gatekeeper for the entry of F-actin in the pellicular space, between the IMC and plasma membrane, to reach other glideosome components, such as myosin A (MyoA), and sustain parasite forward motion [11, 17]

**Fig 1.**
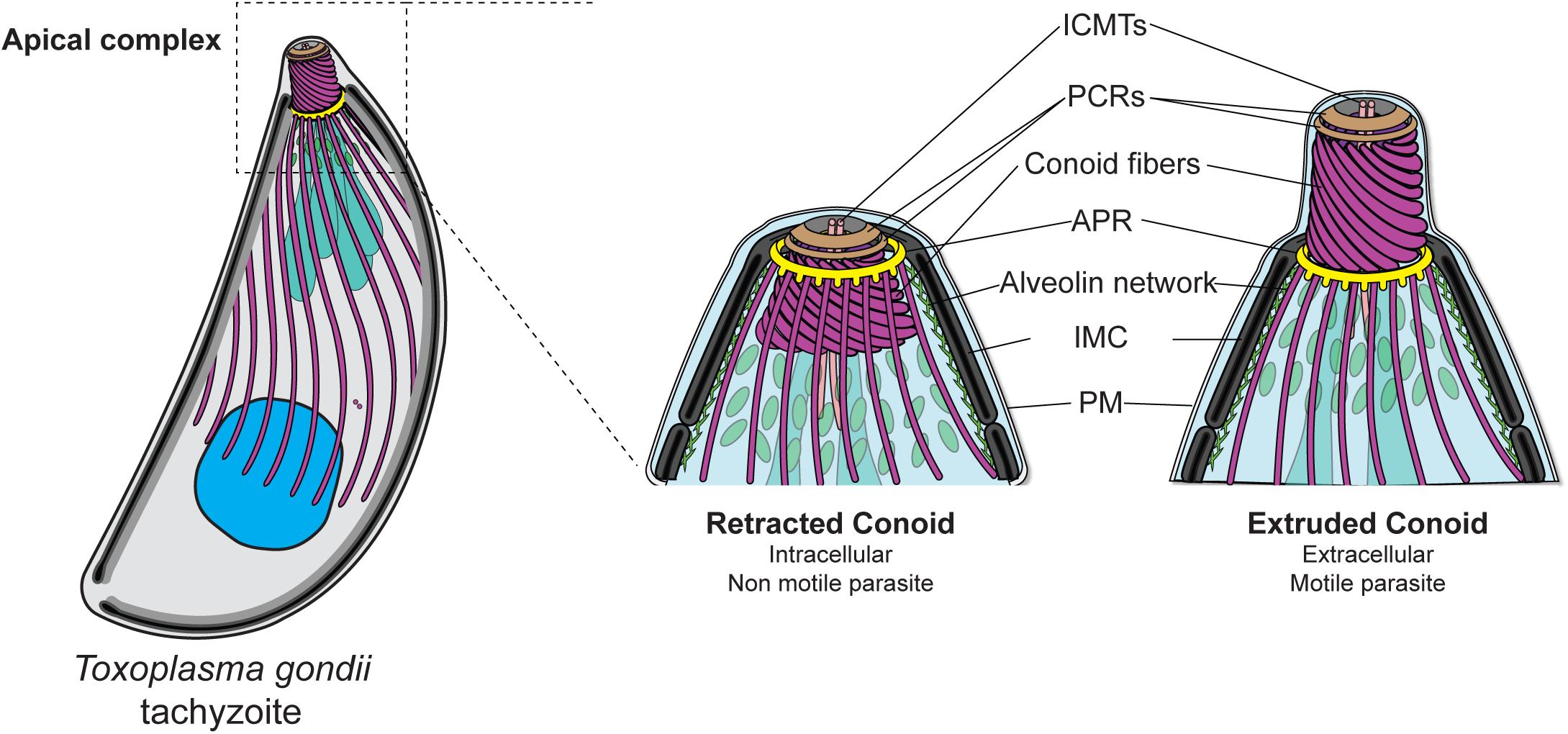
*Toxoplasma gondii* tachyzoite cell organization. **Left panel:** general cell organization of *Toxoplasma gondii* tachyzoites. **Right panel:** Zoomed view of the detailed apical complex. ICMTs: Intraconoidal microtubules. PCRs: Preconoidal rings. APR: Apical Polar Ring. IMC: Inner Membrane Complex. PM: Plasma membrane. Scheme modified from [3]

The APR is assumed to act as an atypical microtubule organizing center (MTOC) for the SPMTs [8, 9]. Several mutants that lead to the loss of APR result in disorganization of the SPMTs and disappearance of the conoid [7, 18–21]. Only a few APR proteins have been identified to date in *T. gondii*. KinesinA (KinA) is an early marker of APR, whereas APR1 and RNG1 appear later during daughter cell formation [18, 22]. Individual protein downregulation has a minimal effect on parasite replication; however, simultaneous depletion of KinA and APR1 resulted in the loss of the APR, disorganized SPMTs and drastic defects in gliding motility and invasion [18]. Similarly, depletion of RNG2 was previously reported to alter microneme secretion, leading to motility and invasion defects [23]. Interestingly, RNG2 exhibits an atypical dual localization, with the carboxy-terminus positioned to the APR and the amino-terminus localized to the conoid [23].

The use of recombinantly produced RNG2 shows that the protein is self-processed and prone to oligomerization, supporting a model of APR-conoid tethering that is compatible with rapid extension and shrinking distances. Iterative U-ExM applied to RNG2 and immunoelectron microscopy (IEM) using polyclonal anti-RNG2 antibodies confirmed that this protein spans the gap between the conoid and the APR. Of relevance, 22 binding fibers (BFs) were previously visualized by electron microscopy and postulated to assist conoid extrusion [24]. Conditional depletion of RNG2 results in the dramatic conoid detachment from the apical pole, further supporting its role in tethering the conoid to the APR which is critical retraction to keep the organelle in place

## Results

### RNG2 is self-processed and oligomerizes in vitro

*RNG2* codes for a 290 kDa protein that encompasses a tropomyosin domain (Tropo) within a long coil-coiled region flanked by disordered N- and C-terminal ends (Fig 2A). To decipher the biochemical properties of RNG2, we expressed recombinant full length and truncated RNG2 variants with an N-terminal His-Tag and a C-terminal TwinStrep-Tag in Sf9 insect cells (Fig 2A). All RNG2 constructs showed robust expression; however, only the ΔN+CTer variant exhibited good solubility (Fig 2B). Full-length RNG2 was completely insoluble, whereas the ΔNTer, ΔTropo, and ΔCTer variants showed partial solubility (Fig 2B-C). Western blot analysis with anti-His and anti-Strep antibodies further highlighted the presence of both N- and C-terminally truncated fragments for almost all the variants (Fig 2C). Only the ΔNTer variant did not display any processing, with a single band detected with anti-His and anti-Strep antibodies corresponding to the predicted size of the protein (Fig 2C).

**Fig 2.**
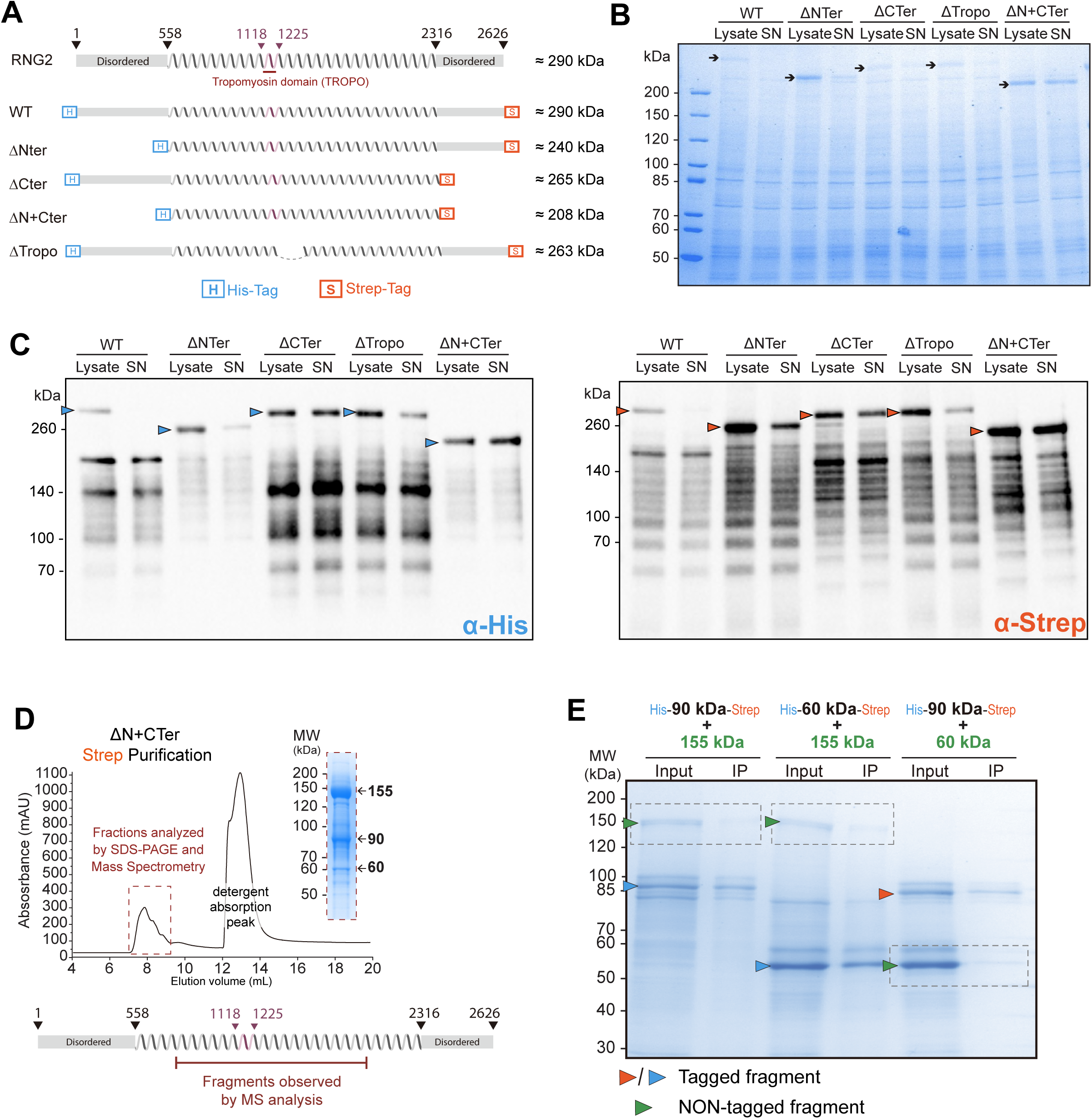
Biochemical investigation of wild-type and processed RNG2. **(A)** Representation of the RNG2 constructs expressed in SF9 insect cells. **(B)** Insect cell protein expression and solubility of the RNG2 truncation variants, as assessed by Coomassie blue staining. **(C)** Protein expression and solubility of RNG2 truncation variants were assessed via western blotting. **(D)** Size-exclusion chromatography (SEC) chromatogram of the affinity-purified RNG2-ΔN+CTer variant obtained via Strep-tactin purification. A Coomassie-stained gel of the protein-containing fractions of the SEC is shown on the right. **(E)** Pull-down assay of RNG2 fragments assessed by Coomassie blue staining.

Purification via the N-terminal His-Tag was not achievable for any of the constructs. In contrast, purification by pulling on the C-terminal Strep-Tag proved successful for RNG2 constructs where the C-terminal 300 residues were truncated (ΔCTer and ΔN+CTer RNG2), moreover requiring salt concentrations largely exceeding physiological conditions (800 mM NaCl). Affinity-purified proteins were subjected to size exclusion chromatography (SEC), and the fractions were analyzed by SDS‒PAGE, mass spectrometry, and western blotting. For both C-terminally truncated constructs, purification led to the isolation of the same three main products with sizes corresponding to 155 kDa, 90 kDa, and 60 kDa (Fig 2D, S1A-B Fig). All the fragments coeluted in the void volume of the SEC, corresponding to molecule sizes greater than 600 kDa, indicating that the fragments formed large oligomers (Fig 2D, S1A-B Fig).

Western blot analysis confirmed that all the purified products had lost the N-terminal His-Tag (S1B Fig). This provides an explanation for the failure to isolate RNG2 fragments by using the N-terminal His tag and strongly suggests that these fragments are self-processed. Mass spectrometry (MS) analysis confirmed that the purified material corresponded to RNG2 fragments and allowed mapping of the fragment boundaries (Fig 2D, S1C Fig).

To isolate homogeneous RNG2 truncated proteins, each of the 155, 90 and 60 kDa fragments was individually expressed and purified via the same workflow as described earlier. Like the larger constructs, each fragment eluted in the void volume by SEC, indicating oligomerization (S1D Fig). To test whether fragments of different sizes could form hetero-oligomers, pull-down experiments were performed. Different pairs of purified fragments were mixed and pulled down by affinity purification using a tag present only on one of the fragments (Fig 2E). None of the fragments coeluted with a partner, indicating that preformed homo-oligomers cannot assemble into hetero-oligomers (Fig 2E).

Collectively, heterologous expression of RNG2 provided evidence that the protein is highly self-processed. Only a mixture of truncated RNG2 constructs could be purified, which were associated with large homo-oligomers containing fragments of 155, 90 and 60 kDa.

### RNG2 forms 22 tethers between the conoid and the APR

Double epitope tagging at both extremities of RNG2 previously revealed that the N-terminal tag stains the conoid, whereas the C-terminal tag labels the APR [23]. To resolve this conundrum and scrutinize the biochemical properties of RNG2 in *T. gondii*, we introduced epitope tags at the N-terminus (Myc), center (Ty), and C-terminus (HA) along with a C-terminal mAID cassette (Fig. 3A). The endogenous triple epitope-tagged RNG2 clonal line was confirmed by genomic PCR (S2A Fig) and western blot analysis (Fig. 3B). Several RNG2 truncated products of high molecular weight were detected with anti-HA and anti-Ty antibodies. The specificity of those bands was confirmed by their disappearance in parasites treated with auxin (IAA). However, the anti-Myc antibody detected only the full-length version of RNG2, which is consistent with N-terminal processing events (Fig 3B).

**Fig 3.**
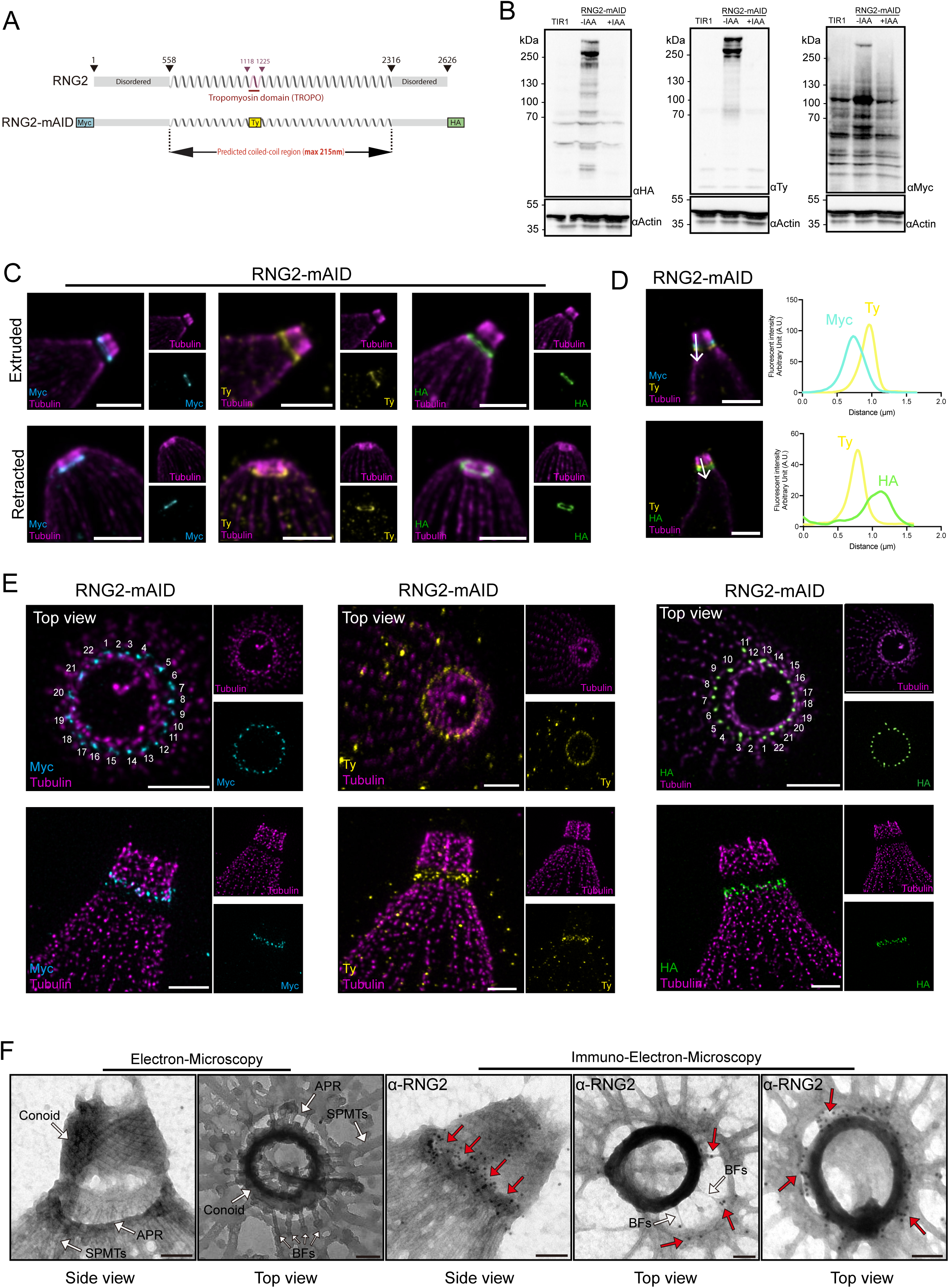
RNG2 spans between the APR and the conoid, forming 22 tethers. **(A)** Illustration of endogenous tagging of RNG2, including three epitope tags and a minimal auxin inducible (mAID) motif. Estimation of the protein length corresponding to the distance coverage of the central part of the protein in linear conformation. **(B)** Protein depletion after 48 h of auxin (IAA) treatment assessed by western blot. **(C)** U-ExM image of the 3 locations of RNG2-mAID depending on the tag insertion. Images were taken with extruded and retracted conoids. Scale bar = 3 μm. **(D)** Colocalization of the Ty internal tag with the N-terminal (Myc) or C-terminal (HA) tags. Scale bar = 3 μm. Plot profile analysis of the signal. **(E)** Iterative expansion microscopy (iU-ExM) of the N-terminal (Myc), middle (Ty) and C-terminal (HA) tags of RNG2-mAID. Scale bar = 3 μm. **(F) Left panel:** Electron microscopy images of the *T. gondii* apical complex as an ultrastructure control. **Right panel:** Immunoelectron microscopy images obtained with the α-RNG2 antibody. Red arrows = gold particles. Scale bar = 100 nm.

Indirect immunofluorescence assay (IFA) using the three antibodies (Myc, Ty, HA) confirmed the apical localization of the RNG2 as well as tight regulation under auxin treatment (S2B Fig). To refine the dynamic behavior of RNG2, the position of the different tags was assessed via U-ExM on extracellular tachyzoites on extruded or retracted conoids (Fig. 3C). The N-terminus of RNG2 (Myc) was found at the base of the conoid both in extruded and retracted states, whereas the C-terminus (HA) was always localized on top of the SPMTs at the APR level, confirming previous observations [23]. The central Ty tag localized between the conoid base and SPMTs, which was further confirmed through dual localization with the N- and C-terminal tags (Fig. 3D). Interestingly, the Ty-Tag colocalized with the N-terminal Myc-Tag only when the conoid was retracted, suggesting a shortening of the protein length in the retracted state (S2C Fig). Taken together, these data confirm the dual localization of RNG2 [23] and suggest that the protein spans the gap between APR and conoid.

To achieve higher resolution, we employed iterative expansion microscopy (iU-ExM), which allows for a 16-fold expansion ratio compared with the 4-fold expansion ratio of U-ExM [25]. With increased resolution, we observed that the RNG2 C-terminus forms 22 puncta that align with the tips of SPMTs, as well as 22 puncta on the conoid looking at the RNG2 N-terminus (Fig. 3E). In parallel, we raised anti-RNG2 polyclonal antibodies in rabbit by injecting the purified RNG2- ΔCTer fragments described earlier (Fig 2D). The anti-RNG2 antibodies showed very high specificity for RNG2 according to both U-ExM and by western blot analysis (S3A-B Fig). In line with our previous observations, the antibodies covering the central region of the protein decorated the space between the conoid and SPMTs (S3A Fig) and between the signals of Myc and HA-Tags (S3C Fig).

Recently, electron microscopy (EM) ultrastructural analysis revealed the existence of binding fibers (BFs) organized in a cartwheel-like conformation and linking the conoid to the SPMTs [24]. To test whether RNG2 could be a component of BFs, we performed immunoelectron microscopy (IEM) with rabbit anti-RNG2 antibodies (Fig. 3F). Although BFs were detected in the cytoskeleton only as an ultrastructure control (Fig. 3F), in the cytoskeletons processed for IEM, many of the fibers disappeared likely because of the numerous washes and incubations with the antibodies. However, a large accumulation of gold labels was found in the space between the conoid and APR and in some BFs still visible in the preparations (Fig. 3F). Taken together, the iU-ExM- and IEM-based evidence for the spread of RNG2 between the APR support the role of RNG2 as a component of the 22 BFs.

### RNG2 critically maintains the conoid attached to the APR during retraction

Considering the biochemical and imaging observations supporting the role of RNG2 in conoid attachment to the APR, we revisited its function, taking advantage of the fast depletion of the protein with the auxin degron system. RNG2 depletion by IAA, as well as full deletion of the gene (RNG2-KO), were confirmed to be fitness conferring by plaque assays (S4A Fig). Conoid extrusion can be triggered by artificial elevation of cGMP levels with BIPPO, a phosphodiesterase (PDE) inhibitor [26] and unambiguously observed by U-ExM and EM [11, 26, 27]. Strikingly in BIPPO-stimulated extracellular RNG2-depleted parasites, the conoid detached from the apex of the cell and was found floating in the parasite cytosol both by U-ExM and EM (Fig. 4A-B). Importantly, conoid detachment was prominently observed in more than 80% of BIPPO-stimulated extracellular parasites. To determine whether actomyosin-driven conoid extrusion triggers conoid detachment, we performed a conoid extrusion assay using parasites pretreated with an inhibitor of actin polymerization, cytochalasin D (CD) [11]. Under normal conditions, BIPPO stimulation resulted in more than 84% of parasites exhibiting conoid extrusion, whereas treatment with CD almost completely inhibited extrusion, with 95% of parasites showing a retracted conoid (Fig. 4C). However, in RNG2-depleted parasites, only 9% of stimulated parasites displayed an extruded conoid, while 77% exhibited conoid detachment. In contrast, when treated with CD, most RNG2-depleted parasites maintained a retracted conoid, like the parental strain (Fig. 4C). These findings suggest that conoid extrusion must be stimulated to observe the detachment caused by RNG2 depletion. Imaging of the detached conoid via EM revealed that the organelle remained intact with PCRs and ICMTs still associated with the tubulin cone (Fig. 4B). Additionally, the electron-dense structure capping the 22 SPMTs like a rubber-band, was unaffected (Fig. 4A). The consistent localization and expression levels of conoid and APR markers, conoid protein hub 1 (CPH1) and APR1 [18, 28], respectively further validated these structural observations (Fig. 4AD). Taken together, these findings demonstrate that RNG2 plays a critical role in retaining the conoid at the apical pole following its extrusion.

**Fig 4.**
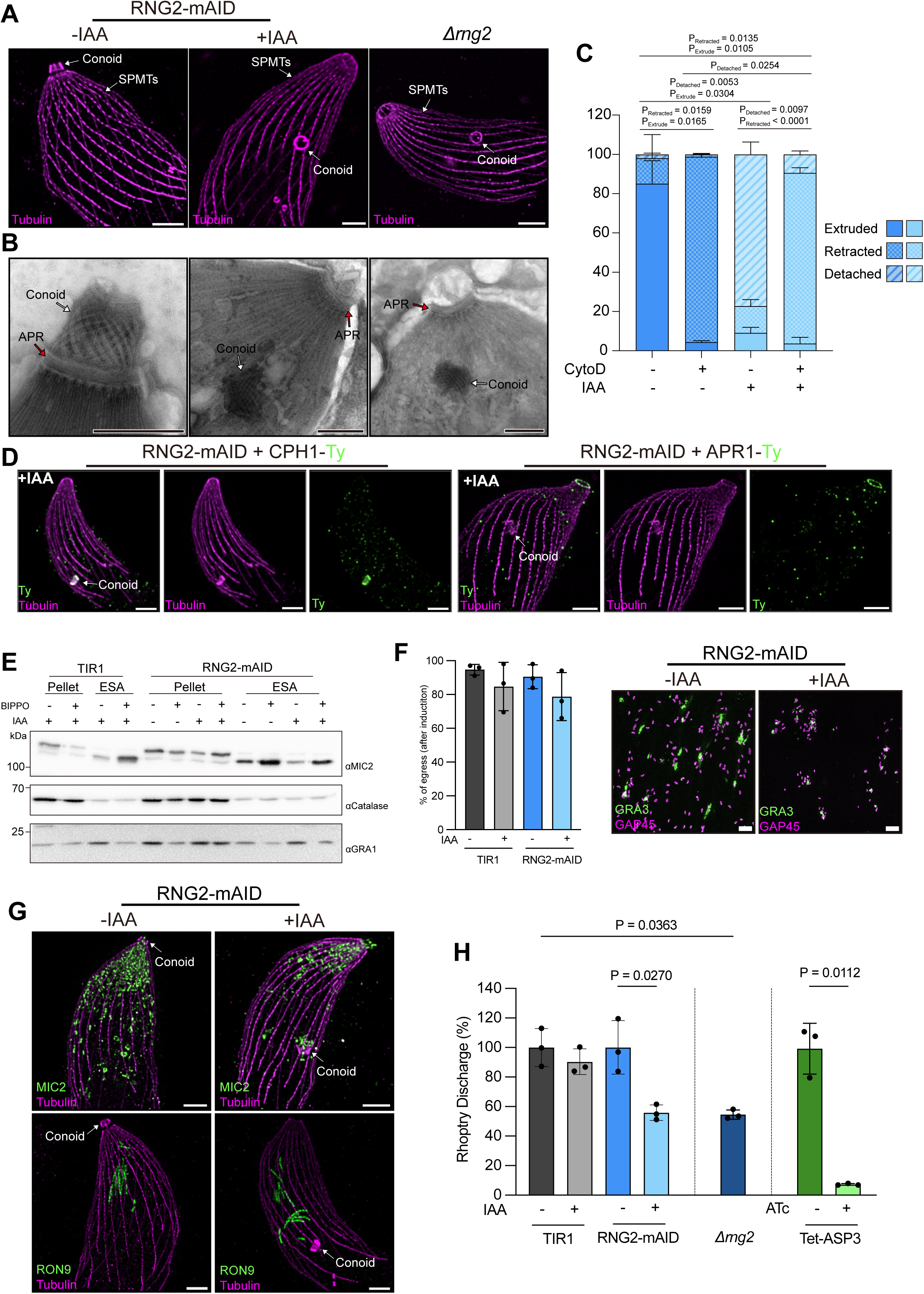
RNG2 depletion leads to conoid detachment and rhoptry mislocalization. **(A)** U-ExM images showing extruded conoid and detached conoid RNG2-mAID-depleted parasites and RNG2-KO parasites. Scale bar = 3 μm. **(B)** Electron microscopy images of negatively stained tachyzoites with extruded conoids and detached conoids in RNG2-mAID- depleted parasites and RNG2-KO parasites. Scale bar = 500 nm. **(C)** Quantification of the conoid state in three categories: extruded, retracted and detached. Under all conditions, extracellular tachyzoites were triggered for conoid extrusion via BIPPO in the absence of auxin (IAA) and with or without cytochalasin D (CD). N = 100 parasites in 3 biological replicates. **(D)** U-ExM images of CPH1 (conoid marker) and APR1 (APR marker) without or without the conoid detached from the APR. Scale bar = 3 μm. **(E)** Western blot analysis of BIPPO-triggered microneme secretion. ESA = Excreted secreted antigens. **(F) Left panel:** quantification of lysed parasitophorous vacuoles upon egress triggered by BIPPO. **Right panel:** representative images of the egress assay. Scale bar = 20 μm. **(G)** U-ExM of secretory organelles, with MIC2 used as a microneme marker and RON9 used as a rhoptry neck marker. Images taken with the conoid attached or detached from the APR. Scale bar = 3 μm. **(H)** Quantification of rhoptry secretion was performed to assess STAT-6 phosphorylation. At least 200 cell nuclei were counted for data plotting, n = 3. ATc = anhydrotetracycline.

### Detachment of the conoid in absence of RNG2 impairs motility and positioning of the rhoptries

*T. gondii* tachyzoites possess eight to twelve rhoptries, with two of them having their necks docked inside the conoid and primed for exocytosis (Mageswaran et al., 2021). Micronemes similarly accumulate at the apical pole of the parasite (Dubois and Soldati-Favre, 2019). Since the conoid is assumed to serve as funnel for the secretion of microneme and rhoptry contents, we assessed the impact of conoid detachment on these processes. Induced microneme secretion showed no significant defect following conditional depletion of RNG2 (Fig. 4E). Consistent with this observation, RNG2-depleted parasites were able to successfully egress, lysing both the parasitophorous vacuole membrane (PVM) and the plasma membrane of the infected host cell upon BIPPO stimulation (Fig. 4F). However, RNG2-depleted parasites remained clustered at the site of host cell lysis, indicating an impairment in motility and dissemination. This was confirmed by a severe defect in the gliding trail assay (S4B Fig). Interestingly, U-ExM revealed that some micronemes were still associated to the detached conoid, pointing to the existence of physical interactions between the conoid and these organelles (Fig. 4G). More strikingly the rhoptries were considerably disorganized in RNG2-depleted parasites (Fig. 4G). Given that proper rhoptry positioning is crucial for their exocytosis, we assessed the ability of the parasites to discharge their rhoptry contents into the host cell in the absence of RNG2. Rhoptry secretion was measured by monitoring the phosphorylation of host cell nuclear STAT-6, a marker for rhoptry discharge [29]. In ASP3-knockdown parasites, used as a negative control, rhoptry secretion was shown to be entirely blocked. In contrast, RNG2-knockdown parasites exhibited a 50% reduction in rhoptry discharge, which was comparable to that observed in Δrng2 mutant (Fig. 4H) [30]. Consequently, depletion of RNG2 resulted in a reduction of invasion efficiency to less than half of that seen in the parental strain (S4C Fig).

To conclude, the disconnection of the conoid from the APR led to improper localization of the rhoptries, which adversely affected the parasite’s invasion capacity. Although microneme secretion and PVM rupture were not impacted in RNG2-depleted parasites, the detachment of the conoid and its PCRs deprived the parasite of apical F-actin polymerization resulting in a profound motility defect.

### The central region of RNG2 is critical for conoid tethering at the apical pole

The coiled-coil region in the central part of RNG2 expands more than two-thirds of the protein length and contains a putative tropomyosin motif insert (Fig 5A). The N-terminal 500 residues and C-terminal 300 residues of RNG2 have no identifiable structure or motifs. To investigate whether RNG2 subdomains are involved in conoid tethering, the *Δrng2* cell line was complemented with truncated versions of the protein expressed under the control of the tubulin promoter. Western blot analysis confirmed the expression of each variant at the expected size, as well as the presence of processed forms like those observed for the endogenous protein (Fig. 5B). IFA and U-ExM of all the truncated proteins confirmed APR localization, with an additional signal in the cytoplasm likely due to overexpression (Fig. 5C, S5A Fig). Plaque assays were performed to assess the fitness of the complemented lines. The overexpression of the full length, ΔNTer, and ΔTropo variants fully rescued the fitness of parasites following the depletion of endogenous RNG2 (Fig. 5D, S5A Fig). However, the ΔCTer and ΔN+CTer variants only partially restored parasite fitness, indicating that the C-terminal region of RNG2 plays a critical role in its physiological function, whereas the N-terminal region and the central tropomyosin domain are dispensable (Fig. 5D, S5A Fig). Consistent with the plaque assay findings, the ΔCTer- and ΔN+CTer-complemented strains exhibited mild invasion defects, whereas the WT and ΔTropo strains fully complemented the initial invasion defect (Fig. 5E). Notably, the ΔNTer variant showed a slight but significant reduction in invasion (10% reduction).

**Fig 5.**
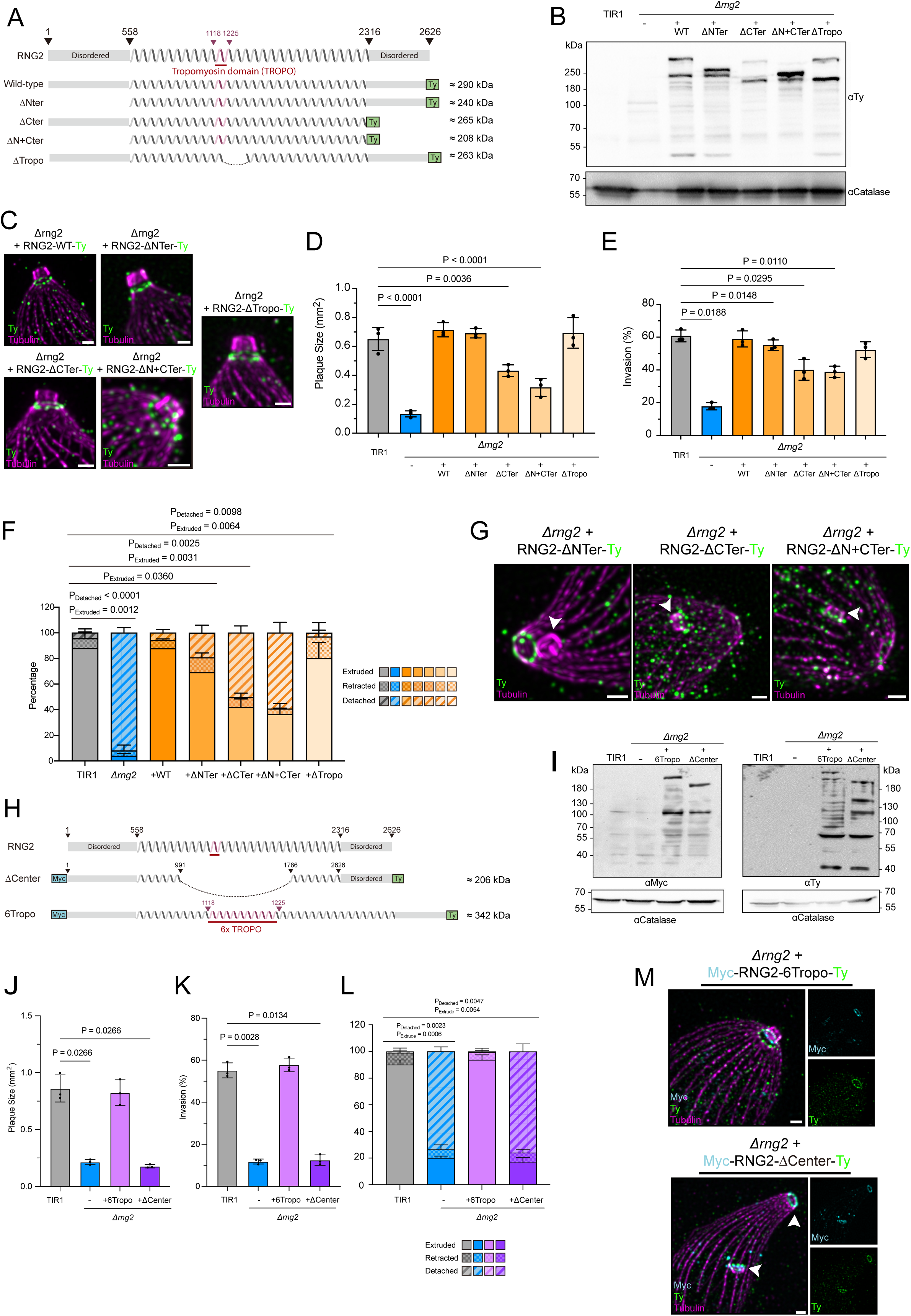
Investigation of RNG2 subdomain functions. **(A)** Representation of the RNG2 variants expressed in the RNG2-KO background. **(B)** Western blot analysis of the expression of RNG2 variants expressed in the RNG2-KO background. **(C)** U-ExM images of the localization of the RNG2 variants. Scale bar = 1 μm. **(D)** Quantification of the plaque sizes created by the different RNG2 strains via a plaque assay. Ten plaques were measured per biological replicate. Three biological replicates were evaluated. **(E)** Quantification of the percentage of invasion by the different RNG2 strains in the invasion assay. One hundred parasites were measured per biological replicate. Three biological replicates were evaluated. **(F)** Quantification of the percentages of conoid retraction, extrusion, and detachment by the different RNG2 strains in the conoid detachment assay. One hundred parasites were measured per biological replicate. Three biological replicates were evaluated. **(G)** U-ExM images of the localization of the RNG2 variants under conditions in which the conoid detached from the apical pole. Scale bar = 1 μm. **(H)** Representation of the second round of RNG2 variants expressed in the RNG2-KO background. **(I)** Western blot analysis of the expression of the second round of RNG2 variants expressed in the RNG2-KO background. **(J)** Quantification of the plaque sizes created by second-round RNG2 variants via a plaque assay. Ten plaques were measured per biological replicate. Three biological replicates were evaluated. **(K)** Quantification of the percentage of invading cells in response to the second round of RNG2 overexpression via an invasion assay. One hundred parasites were measured per biological replicate. Three biological replicates were evaluated. **(F)** Quantification of the percentage of conoid retraction/extruded/detaching by the second round of the RNG2 variant in the conoid detachment assay. One hundred parasites were measured per biological replicate. Three biological replicates were evaluated. **(M)** U-ExM images of the localization of the second round of RNG2 variants under conditions in which the conoid detached from the apical pole. Scale bar = 1 μm.

Examination of conoid detachment in the various RNG2 variants revealed that in the Δrng2 strain, over 90% of the conoids were detached from the APR, while in the TIR1 parental strain, fewer than 3% of conoids were detached, possibly due to artifacts from U-ExM (Fig. 5F). The wild-type and ΔTropo variants successfully complemented the depletion of endogenous RNG2, resulting in very few detached conoids. In contrast, 50% of conoids in the ΔCTer and ΔN+CTer variant-complemented strains were detached from the APR (Fig. 5F). Although the ΔNTer strain did not show a significant fitness defect, 20% of its conoids were detached, which corresponds with the slightly impaired invasion phenotype observed (Fig. 5F). Following conoid extrusion, the protein variants either remained tethered to the APR (as seen with ΔNTer) or were associated with the floating conoids (as observed with the ΔCTer variant or in strains with both N- and C-terminal truncations). This suggests that the N-terminal region of RNG2 is associated with the detached conoid, while the C-terminal region interacts with the APR (Fig. 5G).

Having established that the N- and C-terminal extremities of RNG2 are required to bind with the conoid and APR, respectively, we explored the internal sequence that might mediate oligomerization. To this end, two additional RNG2 variants were engineered: one featuring an extended central region with six tropomyosin domain repeats (RNG2-6Tropo) and another shorter variant where most of the internal sequence was removed (RNG2-ΔCenter) (Fig. 5H). Both variants were dually tagged with Myc-tags and Ty-Tags at their N- and C-terminal ends, respectively. Western blot analysis confirmed that both variants were expressed correctly and exhibited several processed forms, particularly in the extended RNG2 protein (Fig. 5I). Comparison of the blots indicated the presence of several fragments of identical size between the two variants, suggesting that a major N-terminal fragment of approximately 110 kDa and two key C-terminal fragments of 60 and 40 kDa were produced (Fig. 5I). These fragments likely correspond to the minimal regions essential for binding to the conoid and APR, respectively. Both variants accumulated at the apical pole, as shown by IFA (S5C Fig). While the 6Tropo variant fully complemented the *Δrng2* strain, the ΔCenter strain failed to complement it leading to conoid detachment (Fig. 5J-K-L, S5B Fig). The 6Tropo variant dually localized to the conoid and APR, likel RNG2 (Fig. 5M). In contrast, RNG2-ΔCenter localized at both the conoid and APR when detected with antibodies recognizing either the N- or C-terminal extremities, suggesting that the mutant protein is too short to span the gap between the two structures (Fig. 5M). These observations establish that i) RNG2-ΔCenter is not processed, leading to colocalization of the N- and C-termini; ii) The extremities of RNG2 are sufficient for binding the conoid and APR but not for conoid tethering; and iii) The processing and oligomerization of the central region of RNG2 are critical for tethering the conoid at the apical pole.

## Discussion

The conoid serves as a critical hub for assembling all the molecular machinery necessary for parasite invasion. This dynamic cytoskeletal element orchestrates the discharge of secretory organelles and the assembly of the actomyosin-based glideosome, which powers parasite motility. Originally believed to be exclusive to cyst-forming parasites, the conoid has recently been found in a broader range of Apicomplexans, including *Plasmodium* and *Cryptosporidium* species [31–34].

In *T. gondii*, the conoid undergoes calcium-dependent extrusion and retraction through the APR, a process crucial for controlling parasite motility by directing actin filaments into the pellicular space, leading to myosin A-driven forward motion [11]. Despite this understanding, the factors responsible for maintaining the conoid at the apical pole and facilitating its movement through the APR remained unclear until recently. The identification of BFs acting as flexible tethers between the APR and conoid provided the first physical link between these structures [24].

RNG2 emerged as a strong candidate for APR attachment due to its dual localization between the APR and conoid [23]. *In vitro* studies revealed that RNG2 is prone to self-processing and oligomerization, with fragments localizing between the conoid and APR. The purified proteins used to generate the RNG2 rabbit polyclonal antibody, which specifically recognizes the processed products, were instrumental in leading us to conclude that RNG2 undergoes processing within the parasites. This processing results in certain RNG2 fragments localizing between the conoid and the APR. High-resolution imaging based on iU-ExM detected RNG2 as 22 puncta at the APR and conoid levels supporting its role as part of the BFs. Of relevance, RNG2 depletion caused conoid detachment upon BIPPO stimulation, emphasizing its role in anchoring the conoid. Notably, this detachment occurs after extrusion, as CD prevents detachment, indicating that RNG2 is crucial for maintaining conoid positioning during and after extrusion.

Microneme secretion remained unaffected in RNG2-depleted parasites, consistent with previous observations that conoid extrusion is not required for secretion [11]. However, some micronemes remained linked to the detached conoid, suggesting intrinsic microneme-binding properties within the conoid. Three major regulators of microneme apical positioning and exocytosis namely, HOOK, FTS and HIP, were previously shown to accumulate at the apical pole [35]. While their precise localization within the apical pole remains elusive, investigations of RNG2-depleted parasites could shed light on their ability to reach the conoid and possibly explain how micronemes remains associated with the detached conoid.

The proper positioning of rhoptries, essential for their discharge, was disrupted by RNG2 depletion, explaining the significant reduction in rhoptry discharge observed in these parasites. Recent studies have demonstrated that during rhoptry discharge, two rhoptries dock at the apical vesicle (AV) located at the parasite membrane [36]. Given the proximity of the AV to the PCRs that detach from the conoid, investigating the positioning of the AV upon conoid detachment would be worthwhile.

Interestingly, RNG2 depletion did not abrogate the parasite’s fitness entirely, with only a 50% reduction in invasion observed. Despite normal microneme secretion and conoid extrusion, RNG2-depleted parasites exhibited motility defects due to conoid detachment, preventing F-actin translocation necessary for motility.

The critical role of the central region of RNG2 in tethering the conoid at the apical pole was highlighted by the similarity in phenotypes between RNG2-ΔCenter constructs and RNG2-KO parasites. The colocalization of RNG2-ΔCenter at both the conoid and APR, when antibodies targeted either the N- or C-terminal regions, indicated that RNG2’s self-processing is necessary for proper tethering. However, the mechanism by which the N- and C-terminal regions of RNG2 anchor themselves to the APR and conoid remains unclear. It is possible that RNG2 interacts directly with tubulin proteins, or alternatively, it may require interactions with specific surface proteins on both the conoid and APR to facilitate this connection.

Based on localization and phenotypic data, a model is proposed where RNG2 maintains the conoid at the apical pole through strong interactions between its N- and C-terminal binding to conoid and APR, respectively (Fig. 6A-B). RNG2 self-processing generates small fragments of its central region that form filaments to span flexibly the distance between the conoid and APR. Notably, the depletion of APR1, which also leads to conoid detachment, suggests a possible interaction between RNG2 and APR1, warranting further investigation (Leung et al. 2017).

**Fig 6.**
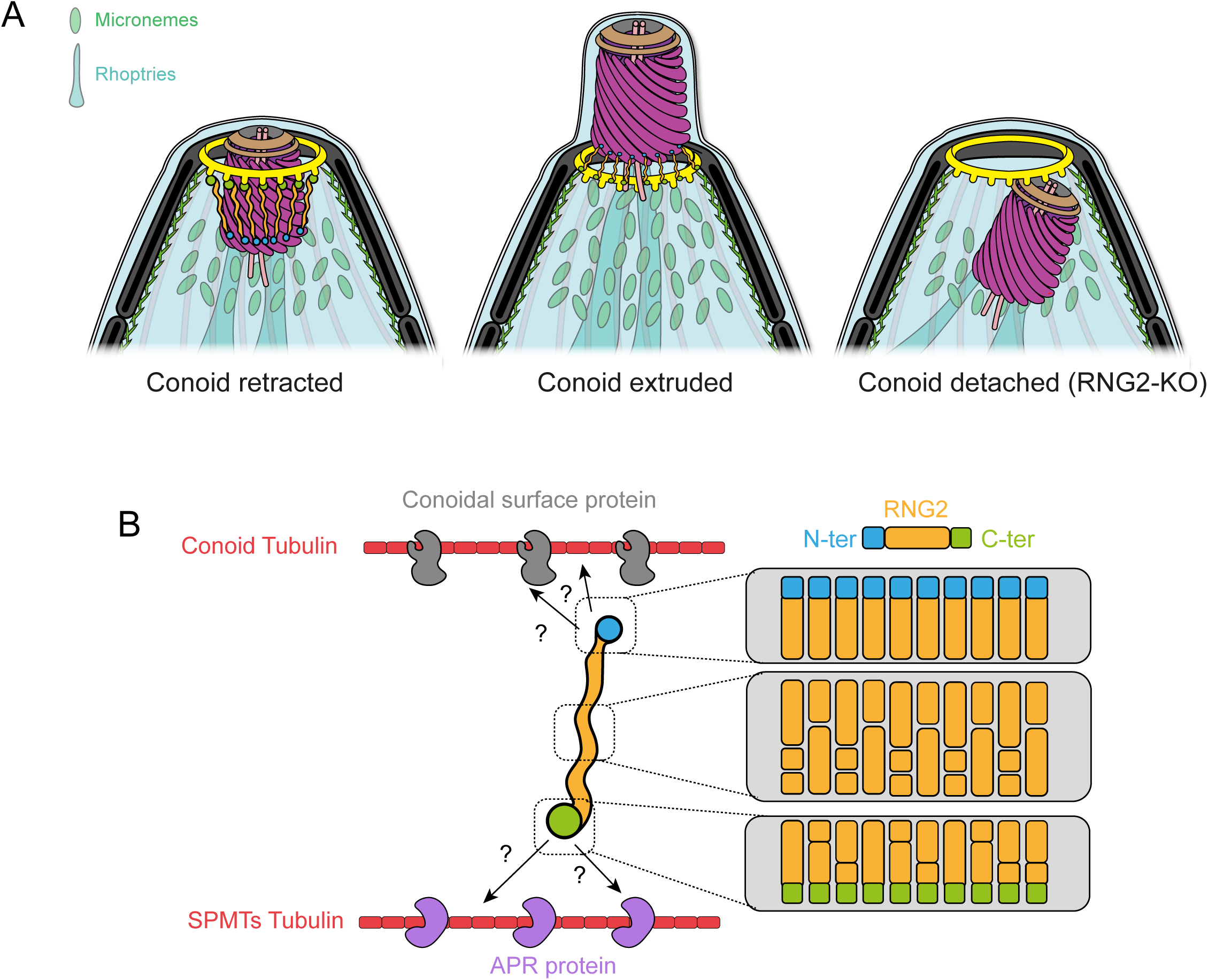
Proposed model for the role of RNG2 and conoid apical anchorage. **(A)** Illustration of the consequences of RNG2 depletion on the apical complex structure of *T. gondii* tachyzoites. **(B)** Illustration of the RNG2 molecular tethering model.

Through a combination of biochemical assays, genetic manipulations, and cell imaging, the pivotal role of RNG2 in connecting the conoid to the APR, while enabling dynamic transitions between extrusion and retraction has begun to be resolved.

## Materials and methods

### Parasite maintenance

*T. gondii* tachyzoites were amplified from HFFs (ATCC) in Dulbecco’s modified Eagle’s medium (DMEM, Gibco) supplemented with 5% fetal calf serum (FCS, Gibco), 2 mM glutamine and 25 µg/ml gentamicin (Gibco). Parasites and HFFs were maintained at 37°C with 5% CO_2_.

### Generation of transgenic cell lines

#### a – Generation of RNG2-mAID (triple epitope-tagged strain)

The gene map of RNG2 was retrieved from the ToxoDB website [37] (www.toxodb.org) via the accession number TGGT1_244470. A specific gRNA (guideRNA) was designed to target the 3’UTR of the endogenous locus via the EuPaGDT tool (www.grna.ctegd.uga.edu) [38]. The sequence of the gRNA (AAGAGAAATTGCCTTCATGT) was inserted into pU6- Universal (pU6-Universal was a gift from Sebastian Lourido, Addgene plasmid #52694) [39]. A repair fragment containing the mAID-HA cassette, and flanking regions was amplified via PCR via KOD polymerase (Novagen, Merck) via the oligos and template listed in Table S1. Freshly egressed RHΔKu80ΔTir1 (called Tir1) tachyzoites were transfected via electroporation [40] with 40 µg pU6-Universal bearing the gRNA alongside 100 µL (2 PCRs of 50 µL) of repair fragment containing the mAID-HA cassette and homology regions for the gene. For enrichment of the transfected population, parasites carrying an HXGPRT cassette were selected with 25 mg/ml mycophenolic acid (MPA) and 50 mg/ml xanthine. The parasites were then cloned and inserted into p96w, and the clonality of the population was assessed via integration PCR, as shown in S2A Fig.

To insert a 3Myc tag on the N-terminal side of the RNG2 gene, two gRNAs were designed to target the 5’UTR of the gene (CAACCGTTCCATGTAAGGCC) and a few base pairs after the START codon (TTCGGGAGACGTTTCTCCTA). The two gRNAs were inserted into the NdeI linearized plasmid listed **in Table S1** via Gibson assembly. A repair fragment of approximately 500 bases containing the 3-Myc-Tag inserted just after the start codon and containing homology regions on each side was synthesized (g-block IDT DNA company). This repair fragment was inserted into the pCR-Blunt-II-Topo vector (Thermo Fisher, K270020) and then amplified via PCR via KOD polymerase in the same way as described previously.

Freshly egressed clonal parasites bearing the mAID-HA cassette were transfected via electroporation as described previously and sorted via flow cytometry (Cas9-GFP). The selected parasites were then cloned and inserted into p96w, and the clonality of the population was assessed via PCR, as shown in S2A Fig.

To insert a 3Ty-Tag in the internal sequence of the RNG2 gene, a gRNA was designed to target this specific region (GAAGGGCAGAAGAACTT) and was inserted into the pU6-universal vector as described above. A repair fragment of approximately 500 bases containing the 3-Ty-Tag inserted just after the in frame of the sequence and containing homology regions on each side was synthesized (g-block IDT DNA company). This repair fragment was inserted into the pCR-Blunt-II-Topo vector (Thermo Fisher, K270020) and then amplified via PCR via KOD polymerase in the same way as described previously. Freshly egressed clonal parasites bearing the N-Ter Myc-tag and mAID-HA cassette were transfected via electroporation as described previously and sorted via flow cytometry (Cas9-GFP). The selected parasites were then cloned and inserted into p96w, and the clonality of the population was assessed via PCR, as shown in For all the assays involving AID-based conditional knockdown systems, protein depletion was achieved by adding 500 µM auxin (IAA) [41].

#### b– Generation of the RNG2-KO strain

To generate the RNG2 knockout line, two gRNAs were designed to target the 5’UTR of the gene (CAACCGTTCCATGTAAGGCC) and the 3’UTR of the gene (TGTGTGTTCATACTCCCGTG). The two gRNAs were inserted into the NdeI linearized plasmid listed in **supplementary table 1** via Gibson assembly. A repair fragment containing the DHFR cassette as well as flanking regions on each side was amplified via PCR via KOD polymerase. Freshly egressed RHΔKu80ΔTir1 (called Tir1) parasites were transfected via electroporation as described previously. For enrichment of the transfected population, parasites carrying the DHFR cassette were selected with 1 µg/ml pyrimethamine.

The selected parasites were then cloned and inserted into p96w, and the clonality of the population was assessed via PCR, as shown in S2A Fig.

#### c– RNG2 variant expression at the UPRT locus

The variant wild-type RNG2 gene was synthesized into the pFastBac vector between the BamHI and HindIII cloning sites (Genscript Company). The RNG2 gene was then subcloned and inserted into the pTub-UPRT-G13-Ty vector between the EcoRI and AvrII restriction sites and under the control of the tubulin promoter to generate the vector pTub-UPRT-RNG2-WT- Ty [42]. For each of the RNG2 variants, specific restriction sites were placed alongside the RNG2 gene sequence, allowing the insertion/deletion of the desired sequences.

For RNG2-ΔNter, pTub-UPRT-RNG2-WT-Ty was digested by the restriction enzyme EcoRI + SpeI and ligated with the primer pairs listed in Table S1 to generate pUPRT-RNG2-ΔNter-Ty.

For RNG2-ΔCter, pFastBac-RNG2-WT was digested by the restriction enzyme NotI to generate pFastBac-RNG2-ΔCter. pFastBac-RNG2-ΔCter was then digested with EcoRI + XbaI and cloned and inserted into pTub-UPRT-G13-Ty, which was subsequently digested with EcoRI+AvrII to generate pTub-UPRT-RNG2-ΔCter-Ty. For RNG2-ΔN+Cter, pTub-UPRT- RNG2-ΔCter-Ty was digested by the restriction enzyme EcoRI + SpeI and ligated with primer pairs 10629/10630 to generate pTub-UPRT-RNG2-ΔN+Cter-Ty. For RNG2-ΔTropo, pFastBac- RNG2-WT was digested by the restriction enzymes SalI + XhoI followed by religation. The sequences were then digested by EcoRI + XbaI to clone into the EcoRI + AvrII digested vector pUPRT-UPRT-G13-Ty to generate pTub-UPRT-RNG2-ΔTropo-Ty.

For RNG2-ΔCenter, the RNG2-ΔCenter variant was synthesized and cloned and inserted into the pTWIST vector. Cloning was performed via the digestion of pTWIST-RNG2-ΔCenter by EcoRI + HindIII and cloning into the vector pTub-MyoL-3Ty, which was digested by EcoRI + HindIII to generate pTub-UPRT-RNG2-ΔCenter.

For RNG2-6Tropo, 6xTropo fragments were synthesized and cloned and inserted into the pTWIST-vector. The 6xTropo fragments were recycled by using the enzymes SalI + XhoI and cloned and inserted into the vector pFastBac-RNG2-WT digested with SalI + XhoI to generate pFastBac-RNG2-6Tropo. pFastBac-RNG2-6Tropo was then digested by EcoRI + XbaI and cloned and inserted into pTub-UPRT-RNG2-ΔCenter digested by MfeI + NheI to generate pTub-UPRT-RNG2-6Tropo

For transfection, *Δrng2* parasites were electroporated with 40 μg of linearized plasmid encoding the selected RNG2 variant together with 20 μg of Cas9-encoding plasmid targeting the UPRT locus [43]. For enrichment of the transfected population, parasites were selected with 5 µM 5-fluorodeoxyuridine (FUDR). The clonality of the parasite lines was assessed via immunofluorescence, and correct expression was assessed via western blotting.

### Expansion microcopy

For this study, the expansion microscopy protocol applied to *Toxoplasma gondii* tachyzoites was followed as previously described [44]. Briefly, parasites resuspended in warm PBS containing 10 μM BIPPO were seeded on 12 mm coverslips precoated with poly-D-lysin (Gibco) for 10 min. After excess PBS was removed, the coverslips were incubated in PBS containing 0.7% formaldehyde and 1% acrylamide for 3 h at 37°C. Polymerization of the expansion gel was performed on ice and contained a monomer mixture (19% sodium acrylate/10% acrylamide/0.1% bis-acrylamide), 0.5% ammonium persulfate (APS) and 0.5% tetramethylenediamine (TEMED). Fully polymerized gels were denatured at 95°C for 90 min in denaturation buffer (200 mM SDS, 200 mM NaCl, 50 mM Tris, pH = 9) and expanded in pure H_2_O overnight.

The next day, the expansion ratio of the fully expanded gels was determined by measuring the diameter of the gels. The well-expanded gels were shrunk in PBS and stained with primary and secondary antibodies diluted in freshly prepared 2% PBS/BSA at 37°C for 2 h. Three washes with PBS/0.1% Tween for 10 min were performed after primary and secondary antibody staining. The stained gels were expanded again in pure H_2_O overnight for further imaging. All U-ExM images used in this study were acquired via Leica TCS SP8 microscope with the lens HC PL Apo 100x/1.40 Oil CS2. Images were taken with Z-stacks and deconvolved with the built-in setting of Leica LAS X. Final images were processed with ImageJ, and the maximum projected images are presented in this study.

### Plaque assay

HFF monolayers were infected with a serial dilution of *T. gondii* tachyzoites and grown for seven days at 37 °C. The cells were fixed with paraformaldehyde-glutaraldehyde for 10 min, followed by neutralization with 0.1 M PBS/glycine. The fixed monolayer was then stained with crystal violet for 2 h and then washed three times with PBS.

### Conoid extrusion assay

To induce conoid extrusion, freshly egressed tachyzoites were incubated with 10 µM BIPPO for 10 min at room temperature. The parasites were then transferred to poly-D-lysine-coated coverslips, followed by an expansion microscopy protocol as described previously. The conoid status of a minimum of 100 parasites was determined for quantification. The data presented in this study were acquired from three biological replicates.

### Invasion assay

Coverslips covered with the HFF monolayer were infected with *T. gondii* tachyzoites, centrifuged at 1200 × g for 1 min and incubated at 37 °C for 30 min. Infected HFFs were fixed with paraformaldehyde-glutaraldehyde and neutralized with PBS/0.1 M glycine. The fixed samples were blocked with 5% PBS-BSA for 20 min at room temperature, followed by 1 h of staining with anti-SAG1 antibody and three washes in PBS. The stained samples were fixed again with 1% formaldehyde for 7 min and permeabilized with PBS/0.2% Triton X-100 for 20 min. The samples were then stained with anti-GAP45 antibody and secondary antibodies. At least 100 parasites per condition were counted to determine the invasion ratio. The data presented in this study were from three biological replicates.

### Microneme secretion assay

Freshly egressed parasites (syringed out in the case of parasites with severe egress defects) were washed twice in warm DMEM, pelleted and resuspended in media containing 10 μM BIPPO. The suspensions were incubated at 37°C for 15 min and centrifuged at 2000 × g to allow separation of the pellet/supernatant (ESA) fraction. The isolated ESA fractions were further centrifuged to remove any remaining cell debris. All the samples were subjected to western blot analysis with anti-MIC2, anti-catalase and anti-GRA1 antibodies.

### Egress Assay

Fresh tachyzoites were used to infect HFF on coverslips, which were subsequently cultured for 30 h at 37 °C. Infected cultures were incubated with DMEM containing BIPPO (10 µM) for 10 min at 37 °C, followed by PFA/Glu fixation and neutralization with 0.1 M PBS/glycine. The coverslips were stained as described previously with the anti-GAP45 antibody and anti-GRA3. At least 100 vacuoles per condition were counted. The data presented in this study were acquired from three biological replicates.

### Gliding trail assay

Freshly egressed tachyzoites were resuspended in warm DMEM containing 10 μM BIPPO, seeded on 12 mm coverslips coated with poly-L-lysine, and centrifuged at 1200 × g for 2 min. The plates with coverslips were incubated for 30 min at 37°C and fixed/stained as described. The anti-SAG1 antibody was resuspended in 2% PBS/BSA, and the corresponding fluorescent secondary antibody was used for visualization by imaging.

### STAT-6 rhoptry secretion assay

A total of 5 × 10^6^ freshly egressed parasites were harvested and resuspended in 250 μl of DMEM for infection of one coverslip seeded with an HFF monolayer. After 30 seconds of centrifugation at 1000 × g, the coverslips were incubated at 37°C for 20 min, followed by 8 min of fixation in ice-cold methanol (MeOH). After 30 min of blocking with 5% PBS/BSA, the coverslips were incubated overnight at 4°C with a STAT6-P antibody. The next day, the coverslips were visualized with a fluorescent secondary antibody and DAPI. The rhoptry discharge efficiency was determined by the ratio of the number of STAT6-P-positive cell nuclei to the total number of nuclei (visualized by DAPI). At least 200 cell nuclei were counted for quantification. The data presented in this study were acquired from three biological replicates.

### Baculovirus-infected Insect cell expression and purification

All recombinant RNG2 proteins were expressed in Sf9 insect cells and purified following a similar protocol. All the plasmids encoding RNG2 possessed an N-terminal His10-tag followed by a TEV recognition sequence and an HRV-3C recognition sequence followed by a TwinSTREP tag. Baculoviruses were generated following standard procedures, and the cells were infected for 48 h at 27°C in a shaking incubator. Purifications were performed via a combination of STREP-tactin affinity purification and size-exclusion chromatography. Briefly, a pellet from 600 ml of cells was resuspended in 120 ml of lysis buffer (PBS supplemented with 800 mM NaCl, 1 mM EDTA, 0.1% Triton X-100 and 3 mM beta-mercaptoethanol) and lysed by shear force using a LM-20 microfluidizer set at 20,000 psi. The lysate was centrifuged at 35,000 × g for 35 min at 4°C, and the supernatant was applied to a 5 ml STREP-TACTIN XT 4Flow column (IBA) at 0.5 ml/min. The column was washed with 50 ml of lysis buffer and eluted with 15 ml of lysis buffer supplemented with 50 mM biotin. The eluted protein was concentrated to 1 ml via a 30 MWCO Amicon concentrator and loaded on a Superdex 200 10/300 GL column at 22°C previously equilibrated in PBS supplemented with 3 mM DTT. Fractions were analyzed via SDS‒PAGE, pooled and flash frozen in liquid nitrogen before being stored at -80°C.

### Western blot analysis for monitoring auxin-induced protein degradation

HFF monolayers seeded in 6 cm petri dishes were infected with freshly egressed parasites and grown for 48 h in the absence or presence of IAA (500 µM). The parasites were pelleted at 1200 rpm, resuspended in protein loading buffer containing 2% SDS and boiled for 15 min at 95°C. To monitor the depletion over various periods, the HFF monolayer was infected with freshly egressed parasites for 1 h at 37 °C. The extracellular parasites were then washed with DMEM supplemented with 500 μM IAA for 1, 2, 4, 8, or 12 h. After 12 h of depletion, all the samples were pelleted and resuspended in SDS. Protein depletion was assessed by Western blotting using an anti-HA antibody against the protein of interest and anti-MIC2 as a loading control.

### Electron microscopy

Extracellular tachyzoites were prepared in the same manner as previously described [19]. Briefly, the extracellular parasites were pelleted in PBS. Conoid extrusion was induced by incubation with 40 µl of BIPPO in PBS for 5 min at 37 °C. A 4 µl sample was applied to a glow-discharged 200-mesh Cu electron microscopy grid for 10 min. The excess sample was removed by blotting with filter paper and immediately washed three times in double-distilled water. Finally, the sample was negatively stained with a 0.5% aqueous solution of phosphotungstic acid for 20 s and air-dried. Electron micrographs of parasite apical poles were collected with a Tecnai 20 transmission electron microscope (FEI, Netherland) operated at an acceleration voltage of 80 kV and equipped with a side-mounted CCD camera (MegaView III, Olympus Imaging Systems) controlled by iTEM software (Olympus Imaging Systems).

### Immunoelectron microscopy

The isolation of the cytoskeleton from tachyzoites of *T. gondii* was performed as previously reported [24]. Briefly, a suspension of tachyzoites (1×10^6^/ml) suspended in PHEM solution (10 mM HEPES, 10 mM EGTA, 1 mM MgCl_2_, 50 μg/ml N-Tosyl-L-phenylalanine chloromethyl ketone, 50 μg/ml Nα-p-Tosyl-L-lysine chloromethyl ketone, and 17.4 μg/ml phenylmethylsulfonyl fluoride, pH 6.9) was exposed to 0.1% Triton X-100 in PHEM for 1 min at RT. The cytoskeleton fraction was centrifuged at 200,000 × *g* for 15 min at 4°C. Under such conditions, most of the cytoskeletons were integrated into the typical crescent shape of tachyzoites. To isolate the cytoskeletons in a cartwheel-like conformation, tachyzoites were exposed to Triton X-100 solution for 3 minutes. For immune-electron microscopy (IEM), cytoskeletons were deposited on Ni grids covered with a formvar film (Polysciences, Warrington, PA, USA), fixed with 1% PAF in PHEM, blocked with 0.1% BSA, and then incubated for 2 h with anti-RNG2 (1:100 dilution) diluted in PHEM inside a humid chamber at RT. After washing, the samples were incubated for 2 h with IgG goat anti-rabbit IgG antibodies (ZYMED Laboratories Inc., San Francisco, CA) coupled to 10 nm colloidal gold particles diluted in PHEM, washed with PHEM, counterstained with uranyl acetate and examined via TEM. As a negative control, the cytoskeleton was incubated with preimmune serum and then with a secondary antibody coupled to gold particles.

## Acknowledgments

We are grateful to Paul Guichard, Virgnie Hamel and Vincent Louvel for their teaching and advice for iterative expansion microscopy. We thank Monica E Mondragon Castelán and Sirenia González Pozos for their help in processing the IEM. Micrographs were obtained at the Electron Microscopy Facility (LaNSE, CINVESTAV-IPN, Mexico). We thank the team at the Bioimaging Core Facility, François Prodon, Olivier Brun and Nicolas Liaudet, for their technical assistance and imaging analysis. The project is funded by the Swiss National Science Foundation to DSF (310030_215445 and CRSII5_198545).

## Author contributions

D.S.-F., R.H., and B.R. conceptualized the project. D.S.-F., R.H., B.R., and O.V. conceptualized the methodology, and R.H., B.R., O.V., R.V., N.D.S.P., R.M.F. and B.M. performed the investigations. R.H., B.R. and O.V. performed the formal analysis, and R.H. wrote the original draft. D.S.-F., O.V., N.D.S.P., B.R. and R.M.F. reviewed and edited the manuscript. D.S.-F. and R.M.F acquired funding and obtained resources. All the authors contributed to this article and approved the submitted version.

## References

1. Adl SM, Leander BS, Simpson AGB, Archibald JM, Anderson OR, Bass D, et al. Diversity, Nomenclature, and Taxonomy of Protists. Systematic Biology. 2007;56(4):684–9. doi: 10.1080/10635150701494127.

2. Hu K, Johnson J, Florens L, Fraunholz M, Suravajjala S, DiLullo C, et al. Cytoskeletal components of an invasion machine--the apical complex of Toxoplasma gondii. PLoS Pathog. 2006;2(2):e13. Epub 2006/03/07. doi: 10.1371/journal.ppat.0020013. PubMed PMID: 16518471; PubMed Central PMCID: PMCPMC1383488.

3. Dos Santos Pacheco N, Tosetti N, Koreny L, Waller RF, Soldati-Favre D. Evolution, Composition, Assembly, and Function of the Conoid in Apicomplexa. Trends Parasitol. 2020;36(8):688–704. Epub 2020/06/04. doi: 10.1016/j.pt.2020.05.001. PubMed PMID: 32487504.

4. Paredes-Santos TC, de Souza W, Attias M. Dynamics and 3D organization of secretory organelles of Toxoplasma gondii. J Struct Biol. 2012;177(2):420–30. Epub 2011/12/14. doi: 10.1016/j.jsb.2011.11.028. PubMed PMID: 22155668.

5. Frénal K, Dubremetz J-F, Lebrun M, Soldati-Favre D. Gliding motility powers invasion and egress in Apicomplexa. Nature Reviews Microbiology. 2017;15(11):645–60. doi: 10.1038/nrmicro.2017.86.

6. Harding CR, Frischknecht F. The Riveting Cellular Structures of Apicomplexan Parasites. Trends Parasitol. 2020;36(12):979–91. Epub 2020/10/05. doi: 10.1016/j.pt.2020.09.001. PubMed PMID: 33011071.

7. Tosetti N, Dos Santos Pacheco N, Bertiaux E, Maco B, Bournonville L, Hamel V, et al. Essential function of the alveolin network in the subpellicular microtubules and conoid assembly in Toxoplasma gondii. Elife. 2020;9. Epub 2020/05/08. doi: 10.7554/eLife.56635. PubMed PMID: 32379047; PubMed Central PMCID: PMCPMC7228768.

8. Nichols BA, Chiappino ML. Cytoskeleton of Toxoplasma gondii. J Protozool. 1987;34(2):217–26. doi: 10.1111/j.1550-7408.1987.tb03162.x. PubMed PMID: 3585817.

9. Morrissette NS, Sibley LD. Cytoskeleton of apicomplexan parasites. Microbiol Mol Biol Rev. 2002;66(1):21–38; table of contents. Epub 2002/03/05. doi: 10.1128/mmbr.66.1.21-38.2002. PubMed PMID: 11875126; PubMed Central PMCID: PMCPMC120781.

10. Hu K, Roos DS, Murray JM. A novel polymer of tubulin forms the conoid of Toxoplasma gondii. J Cell Biol. 2002;156(6):1039–50. Epub 2002/03/20. doi: 10.1083/jcb.200112086. PubMed PMID: 11901169; PubMed Central PMCID: PMCPMC2173456.

11. Dos Santos Pacheco N, Brusini L, Haase R, Tosetti N, Maco B, Brochet M, et al. Conoid extrusion regulates glideosome assembly to control motility and invasion in Apicomplexa. Nature Microbiology. 2022;7(11):1777–90. doi: 10.1038/s41564-022-01212-x.

12. Segev-Zarko LA, Dahlberg PD, Sun SY, Pelt DM, Kim CY, Egan ES, et al. Cryo-electron tomography with mixed-scale dense neural networks reveals key steps in deployment of Toxoplasma invasion machinery. PNAS Nexus. 2022;1(4):pgac183. Epub 20220904. doi: 10.1093/pnasnexus/pgac183. PubMed PMID: 36329726; PubMed Central PMCID: PMCPMC9615128.

13. Mageswaran SK, Guerin A, Theveny LM, Chen WD, Martinez M, Lebrun M, et al. In situ ultrastructures of two evolutionarily distant apicomplexan rhoptry secretion systems. Nat Commun. 2021;12(1):4983. Epub 20210817. doi: 10.1038/s41467-021-25309-9. PubMed PMID: 34404783; PubMed Central PMCID: PMCPMC8371170.

14. Dos Santos Pacheco N, Tell i Puig A, Guérin A, Martinez M, Maco B, Tosetti N, et al. Sustained rhoptry docking and discharge requires Toxoplasma gondii intraconoidal microtubule-associated proteins. Nature Communications. 2024;15(1):379. doi: 10.1038/s41467-023-44631-y.

15. Mondrago R, Frixione E. Ca2+-Dependence of Conoid Extrusion in Toxoplasma gondii Tachyzoites. Journal of Eukaryotic Microbiology. 1996;43(2):120–7. doi: 10.1111/j.1550-7408.1996.tb04491.x.

16. Del Carmen MG, Mondragon M, Gonzalez S, Mondragon R. Induction and regulation of conoid extrusion in Toxoplasma gondii. Cell Microbiol. 2009;11(6):967–82. Epub 2009/05/07. doi: 10.1111/j.1462-5822.2009.01304.x. PubMed PMID: 19416276.

17. Martinez M, Mageswaran SK, Guérin A, Chen WD, Thompson CP, Chavin S, et al. Origin and arrangement of actin filaments for gliding motility in apicomplexan parasites revealed by cryo-electron tomography. Nat Commun. 2023;14(1):4800. Epub 20230809. doi: 10.1038/s41467-023-40520-6. PubMed PMID: 37558667; PubMed Central PMCID: PMCPMC10412601.

18. Leung JM, He Y, Zhang F, Hwang YC, Nagayasu E, Liu J, et al. Stability and function of a putative microtubule-organizing center in the human parasite Toxoplasma gondii. Mol Biol Cell. 2017;28(10):1361–78. Epub 2017/03/24. doi: 10.1091/mbc.E17-01-0045. PubMed PMID: 28331073; PubMed Central PMCID: PMCPMC5426850.

19. Nicolas Dos Santos Pacheco NT, Aarti Krishnan, Romuald Haase, Bohumil Maco, Catherine Suarez, Bingjian Ren, and Dominique Soldati-Favre. Revisiting the Role of Toxoplasma gondii ERK7 in the Maintenance and Stability of the Apical Complex. Mbio. 2021. doi: 10.1128/mBio.02057-21

20. Back PS, O’Shaughnessy WJ, Moon AS, Dewangan PS, Hu X, Sha J, et al. Ancient MAPK ERK7 is regulated by an unusual inhibitory scaffold required for Toxoplasma apical complex biogenesis. Proc Natl Acad Sci U S A. 2020;117(22):12164–73. Epub 2020/05/16. doi: 10.1073/pnas.1921245117. PubMed PMID: 32409604; PubMed Central PMCID: PMCPMC7275706.

21. Dos Santos Pacheco N, Tosetti N, Krishnan A, Haase R, Maco B, Suarez C, et al. Revisiting the Role of Toxoplasma gondii ERK7 in the Maintenance and Stability of the Apical Complex. mBio. 2021;12(5):e02057–21. doi: 10.1128/mBio.02057-21.

22. Tran JQ, de Leon JC, Li C, Huynh MH, Beatty W, Morrissette NS. RNG1 is a late marker of the apical polar ring in Toxoplasma gondii. Cytoskeleton (Hoboken). 2010;67(9):586–98. Epub 2010/07/27. doi: 10.1002/cm.20469. PubMed PMID: 20658557; PubMed Central PMCID: PMCPMC2998517.

23. Katris NJ, van Dooren GG, McMillan PJ, Hanssen E, Tilley L, Waller RF. The apical complex provides a regulated gateway for secretion of invasion factors in Toxoplasma. PLoS Pathog. 2014;10(4):e1004074. Epub 2014/04/20. doi: 10.1371/journal.ppat.1004074. PubMed PMID: 24743791; PubMed Central PMCID: PMCPMC3990729.

24. Díaz-Martin RD, Sandoval Rodriguez FE, González Pozos S, Gómez de León CT, Mondragón Castelán M, Mondragón Flores R. A comprehensive ultrastructural analysis of the Toxoplasma gondii cytoskeleton. Parasitology Research. 2022;121(7):2065–78. doi: 10.1007/s00436-022-07534-3.

25. Louvel V, Haase R, Mercey O, Laporte MH, Eloy T, Baudrier É, et al. iU-ExM: nanoscopy of organelles and tissues with iterative ultrastructure expansion microscopy. Nature Communications. 2023;14(1):7893. doi: 10.1038/s41467-023-43582-8.

26. Howard BL, Harvey KL, Stewart RJ, Azevedo MF, Crabb BS, Jennings IG, et al. Identification of Potent Phosphodiesterase Inhibitors that Demonstrate Cyclic Nucleotide-Dependent Functions in Apicomplexan Parasites. ACS Chemical Biology. 2015;10(4):1145–54. doi: 10.1021/cb501004q.

27. Dos Santos Pacheco N, Tosetti N, Krishnan A, Haase R, Maco B, Suarez C, et al. Revisiting the Role of Toxoplasma gondii ERK7 in the Maintenance and Stability of the Apical Complex. mBio. 2021;12(5):10.1128/mbio.02057-21. doi: 10.1128/mbio.02057-21.

28. Long S, Anthony B, Drewry LL, Sibley LD. A conserved ankyrin repeat-containing protein regulates conoid stability, motility and cell invasion in Toxoplasma gondii. Nat Commun. 2017;8(1):2236. Epub 2017/12/23. doi: 10.1038/s41467-017-02341-2. PubMed PMID: 29269729; PubMed Central PMCID: PMCPMC5740107.

29. Ong Y-C, Reese ML, Boothroyd JC. Toxoplasma Rhoptry Protein 16 (ROP16) Subverts Host Function by Direct Tyrosine Phosphorylation of STAT6*. Journal of Biological Chemistry. 2010;285(37):28731–40. d

30 Dogga SK, Mukherjee B, Jacot D, Kockmann T, Molino L, Hammoudi P-M, et al. A druggable secretory protein maturase of Toxoplasma essential for invasion and egress. eLife. 2017;6:e27480. doi: 10.7554/eLife.27480.

31. Koreny L, Zeeshan M, Barylyuk K, Tromer EC, van Hooff JJE, Brady D, et al. Molecular characterization of the conoid complex in Toxoplasma reveals its conservation in all apicomplexans, including Plasmodium species. PLoS Biol. 2021;19(3):e3001081. Epub 2021/03/12. doi: 10.1371/journal.pbio.3001081. PubMed PMID: 33705380; PubMed Central PMCID: PMCPMC7951837.

32. Guizetti J, Frischknecht F. Apicomplexans: A conoid ring unites them all. PLOS Biology. 2021;19(3):e3001105. doi: 10.1371/journal.pbio.3001105.

33. Martinez M, Mageswaran SK, Guérin A, Chen WD, Thompson CP, Chavin S, et al. Origin and arrangement of actin filaments for gliding motility in apicomplexan parasites revealed by cryo-electron tomography. Nature Communications. 2023;14(1):4800. doi: 10.1038/s41467-023-40520-6.

34. Guérin A, Strelau KM, Barylyuk K, Wallbank BA, Berry L, Crook OM, et al. Cryptosporidium uses multiple distinct secretory organelles to interact with and modify its host cell. Cell Host & Microbe. 2023;31(4):650–64.e6.

35. Dubois DJ, Chehade S, Marq JB, Venugopal K, Maco B, Puig ATI, et al. Toxoplasma gondii HOOK-FTS-HIP Complex is Critical for Secretory Organelle Discharge during Motility, Invasion, and Egress. mBio. 2023;14(3):e0045823. Epub 20230424. doi: 10.1128/mbio.00458-23. PubMed PMID: 37093045; PubMed Central PMCID: PMCPMC10294612.

36. Martinez M, Chen WD, Cova MM, Molnár P, Mageswaran SK, Guérin A, et al. Rhoptry secretion system structure and priming in Plasmodium falciparum revealed using in situ cryo-electron tomography. Nature Microbiology. 2022;7(8):1230–8. doi: 10.1038/s41564-022-01171-3.

37. Harb OS, Roos DS. ToxoDB: Functional Genomics Resource for Toxoplasma and Related Organisms. In: Tonkin CJ, editor. Toxoplasma gondii: Methods and Protocols. New York, NY: Springer US; 2020. p. 27-47.

38. Peng D, Tarleton R. EuPaGDT: a web tool tailored to design CRISPR guide RNAs for eukaryotic pathogens. Microbial Genomics. 2015;1(4). doi: 10.1099/mgen.0.000033.

39. Sidik SM, Hackett CG, Tran F, Westwood NJ, Lourido S. Efficient Genome Engineering of Toxoplasma gondii Using CRISPR/Cas9. PLOS ONE. 2014;9(6):e100450. doi: 10.1371/journal.pone.0100450.

40. Soldati D, Boothroyd JC. Transient Transfection and Expression in the Obligate Intracellular Parasite *Toxoplasma gondii*. Science. 1993;260(5106):349–52. doi: doi:10.1126/science.8469986.

41. Brown KM, Long S, Sibley LD. Conditional Knockdown of Proteins Using Auxin-inducible Degron (AID) Fusions in Toxoplasma gondii. Bio Protoc. 2018;8(4). Epub 2018/04/13. doi: 10.21769/BioProtoc.2728. PubMed PMID: 29644255; PubMed Central PMCID: PMCPMC5890294.

42. Sheiner L, Demerly JL, Poulsen N, Beatty WL, Lucas O, Behnke MS, et al. A Systematic Screen to Discover and Analyze Apicoplast Proteins Identifies a Conserved and Essential Protein Import Factor. PLOS Pathogens. 2011;7(12):e1002392. doi: 10.1371/journal.ppat.1002392.

43. Donald RG, Roos DS. Insertional mutagenesis and marker rescue in a protozoan parasite: cloning of the uracil phosphoribosyltransferase locus from Toxoplasma gondii. Proc Natl Acad Sci U S A. 1995;92(12):5749–53. doi: 10.1073/pnas.92.12.5749. PubMed PMID: 7777580; PubMed Central PMCID: PMCPMC41774.

44. Dos Santos Pacheco N, Soldati-Favre D. Coupling Auxin-Inducible Degron System with Ultrastructure Expansion Microscopy to Accelerate the Discovery of Gene Function in Toxoplasma gondii. In: de Pablos LM, Sotillo J, editors. Parasite Genomics: Methods and Protocols. New York, NY: Springer US; 2021. p. 121–37.

