## Supplementary Figures for "RNG2 tethers the conoid to the apical polar ring in *Toxoplasma gondii*: a key mechanism in parasite motility and invasion"

### Supplementary materials

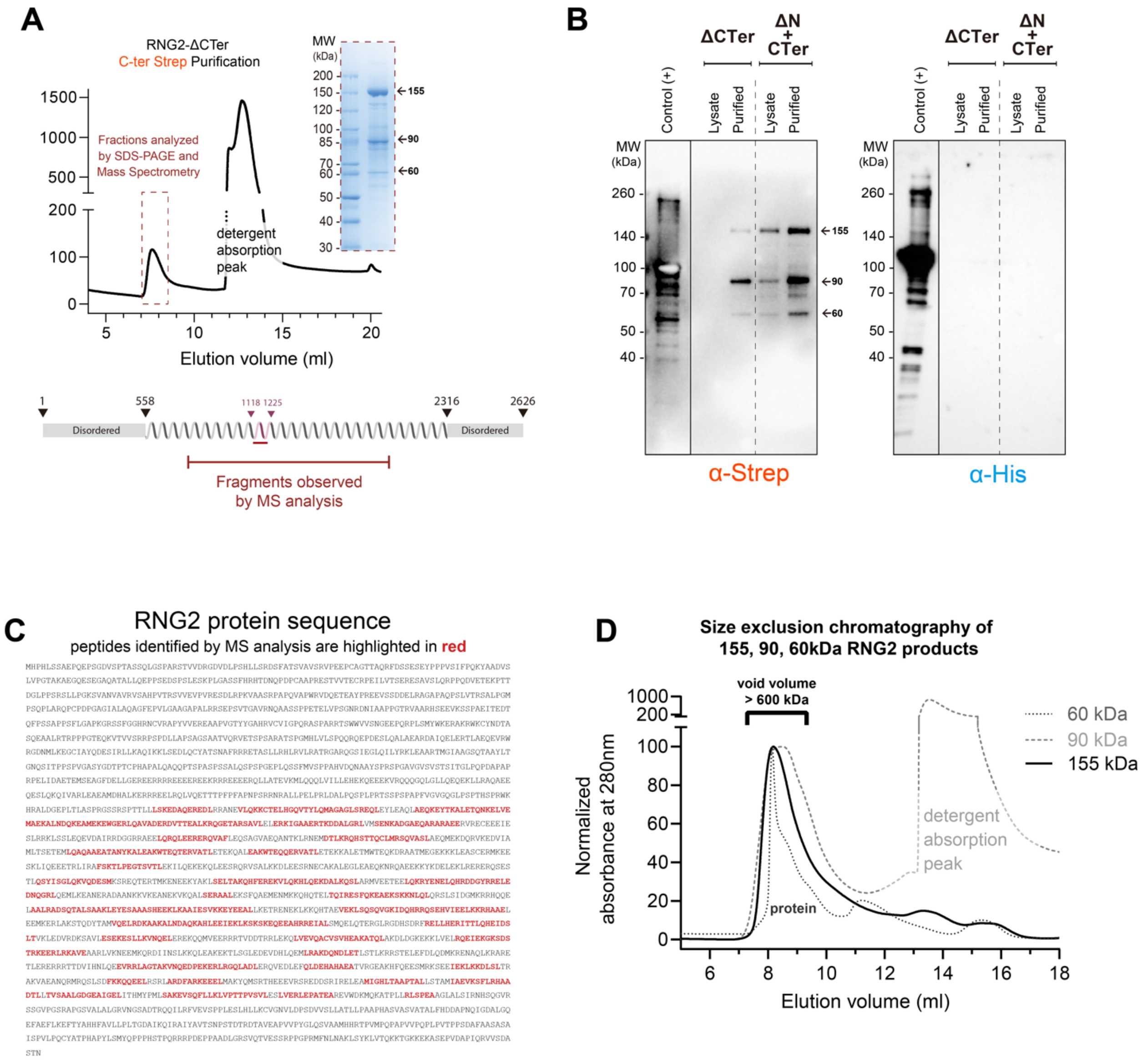


**S1 Fig. RNG2 is self-processed and oligomerizes in vitro**

(**A**) SEC chromatogram of the purified RNG2-ΔCTer variant obtained via Strep-tactin affinity purification, with Coomassie-stained analysis of the protein-containing fractions. **(B)** Western blot analysis of the purification of the ΔN+CTer and ΔCTer variants. **(C)** Mass spectrometry analysis identified peptides belonging to the purified RNG2ΔN+CTer and ΔCTer samples. **(D)** Overlay of the size-exclusion chromatography elution profiles of the 155, 90 and 60 kDa RNG2 fragments. All three fragments similarly eluted in the void volume, corresponding to molecule sizes above 600 kDa.

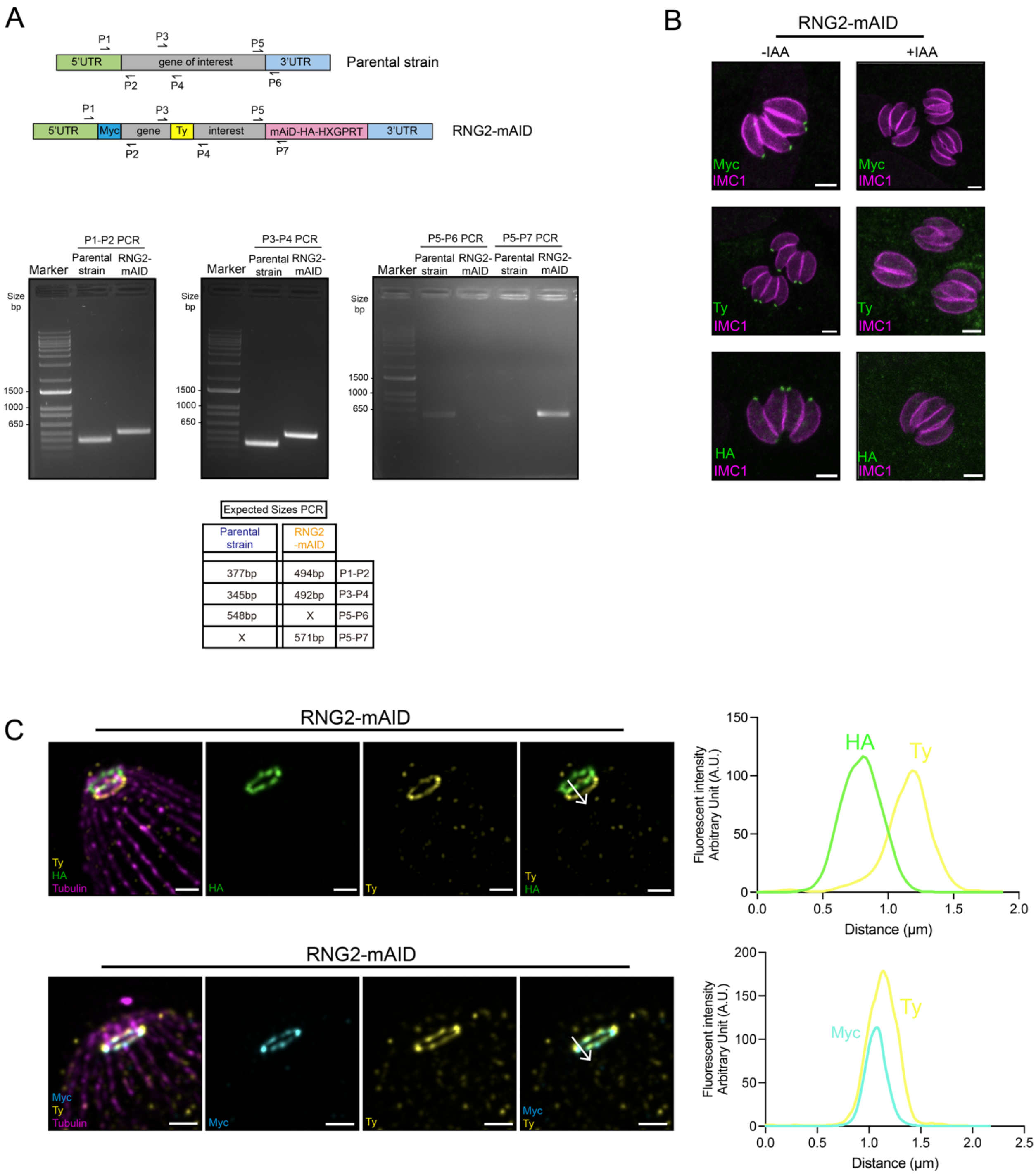


**S2 Fig. Multiple epitopes-tagging across RNG2**

**(A) Upper panel:** schematic representation of the modified endogenous locus. **Lower panel:** polymerase chain reaction of genomic extracted DNA from *T. gondii* tachyzoites to verify the insertion of the N-terminal Myc tag/Internal Ty-Tag and C-terminal mAID-HA. **(B)** Immunofluorescence images of the RNG2-mAID protein in the absence of auxin (12 h) detected via Myc/Ty/HA. Scale bar = 3 μm. **(C)** Colocalization of the Ty internal tag with the N-terminal (Myc) or C-terminal (HA) tags under conoid retraction conditions. Scale bar = 3 μm. Plot profile analysis of the signal.

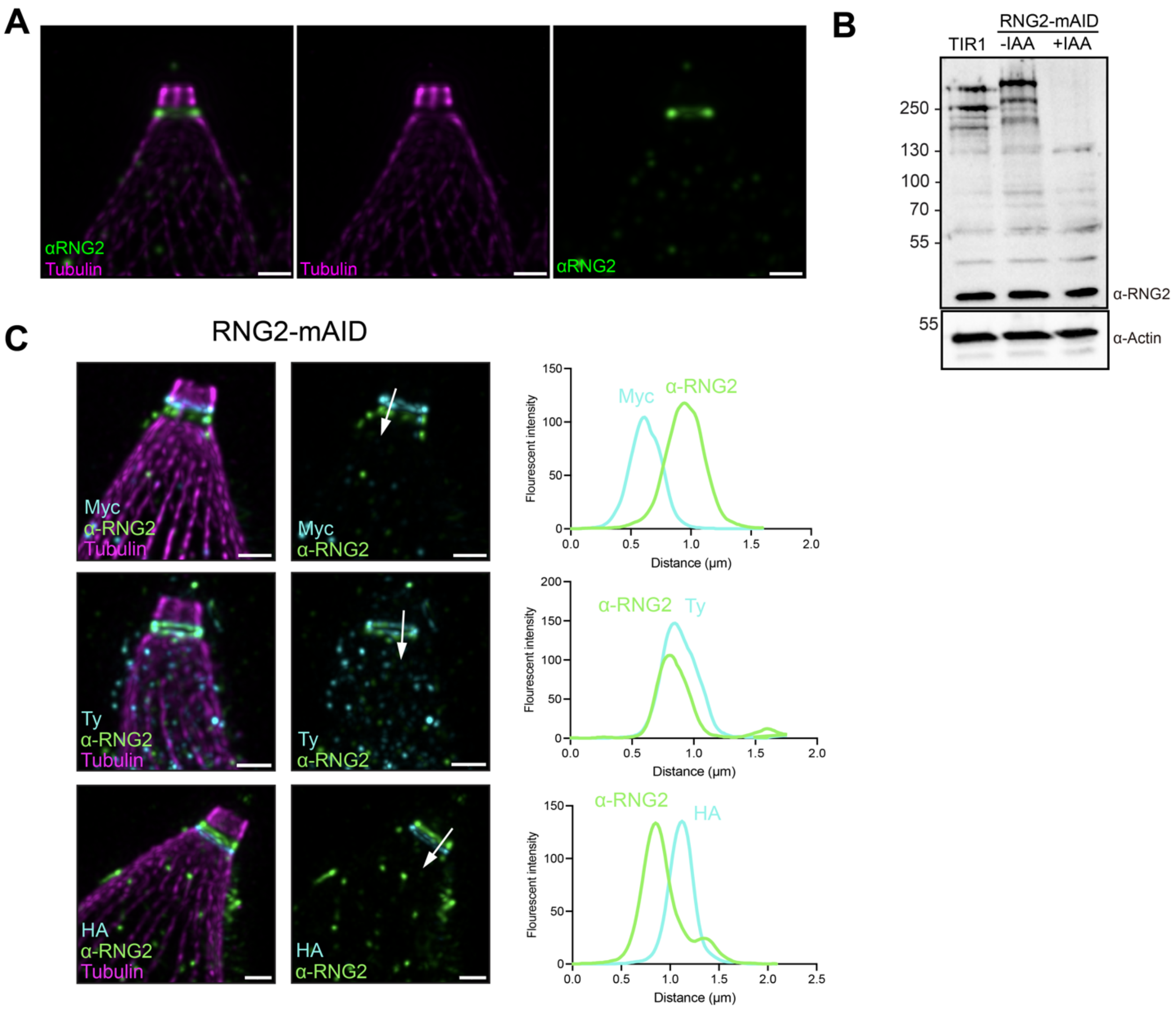


**S3 Fig. RNG2 spans between the APR and the conoid**

**(A)**: U-ExM images of α-RNG2 antibody staining. Scale bar = 1 µm. **(B)**: Western blot analysis using an α-RNG2 antibody in the presence or absence of RNG2. **(C)**: Colocalization of the α-RNG2 antibody with the N-terminal (Myc) or C-terminal (HA) tags. Scale bar = 3 µm. Plot profile analysis of the signal.

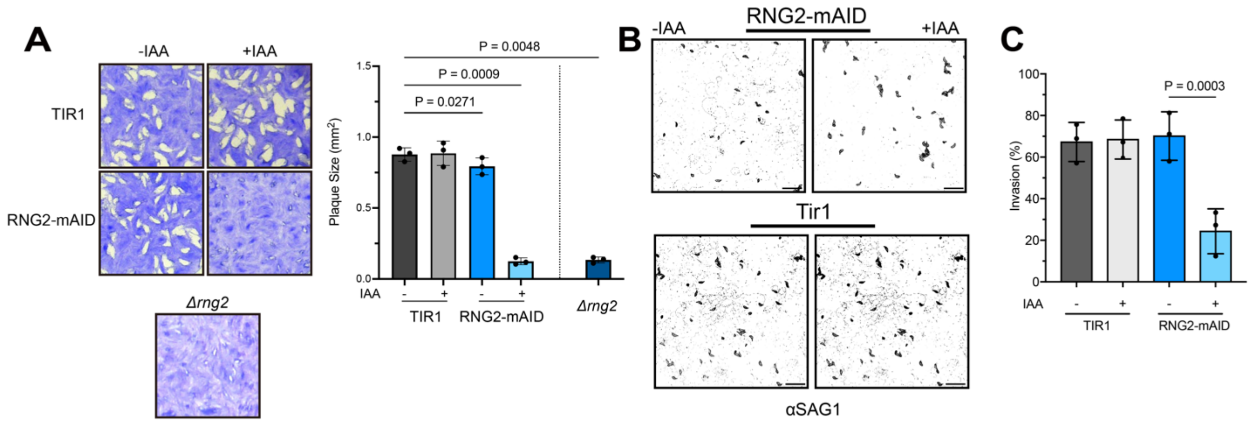


**S4 Fig. Detachment of the conoid in absence of RNG2 impairs parasite motility and invasion** **(A)**: Left panel, images of plaque assays of the RNG2-mAID strain/RNG2-KO strain and TIR1 in the presence or absence of auxin (IAA). **(B)**: Gliding trail assay of RNG2-mAID; gliding motility was determined via visualization of the SAG1 trail after BIPPO induction. **(C)** Quantification of the invasion of the RNG2-mAID strain in the presence or absence of auxin (IAA). One hundred parasites were counted per replicate, and 3 biological replicates were performed. Statistical analysis was performed via one-way ANOVA, *and* $p<0.05$ was considered significant and is plotted in the figure.

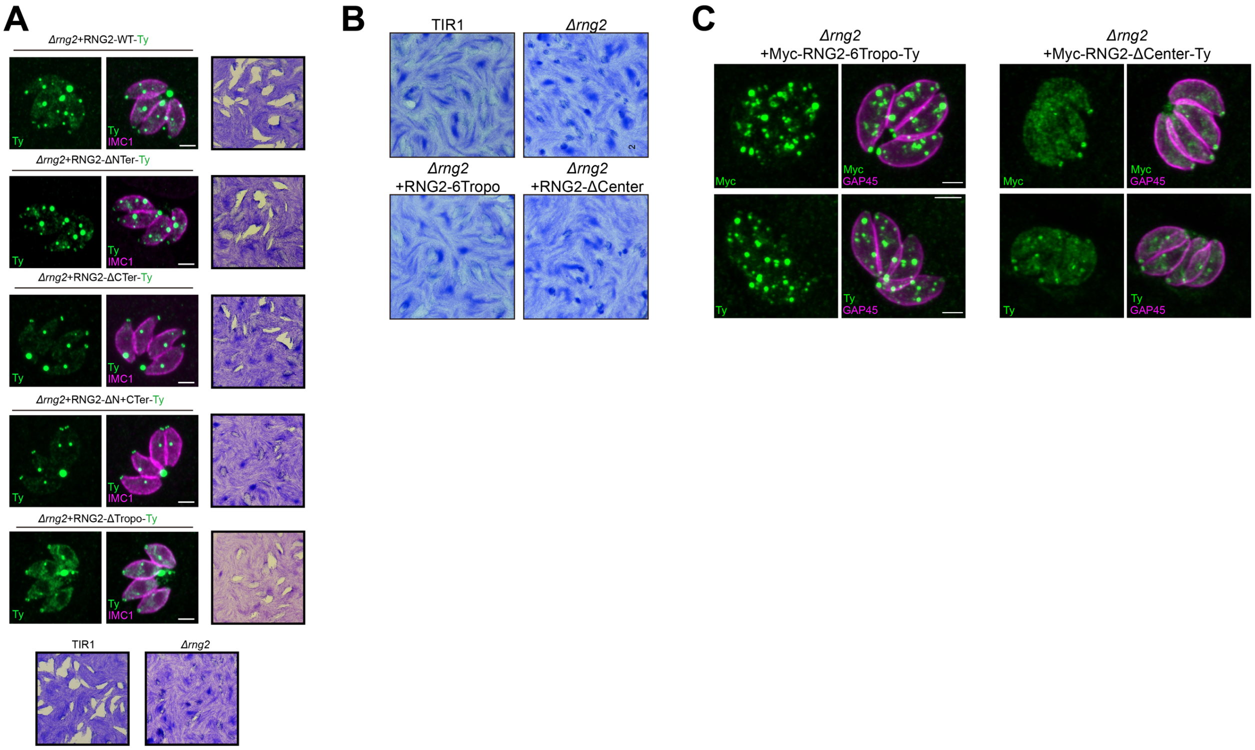


**S5 Fig. functional dissection of the subdomains of RNG2**

**(A)**: IFA localization and plaque assay assessment of the first round of RNG2 variants. Cell marker: IMC1. Scale bar = 2 µm. **(B)**: Plaque assay assessment of the second round of the RNG2 variant. **(C)**: IFA localization of the second round of RNG2 variants. Cell marker: IMC1. Scale bar = 2 µm.
